# Bending the course of evolution: how mutualistic interactions affect macroevolutionary dynamics of diversification in mimetic butterflies

**DOI:** 10.1101/2024.01.26.577270

**Authors:** N. Chazot, M. Pires Braga, T.G. Aubier, V. Llaurens, K. R. Willmott, M. Elias

## Abstract

Evidence that species interactions can affect macroevolutionary dynamics of trait and species diversification is scarce. Mutualistic Müllerian mimicry is a compelling example of example of ecological interactions that has been shown to drive evolutionary convergence, Here, we test how mutualistic Müllerian mimicry shapes macroevolutionary patterns of diversification in the Ithomiini butterflies. We show that the age of color patterns is the primary determinant of species richness within mimicry rings but not phylogenetic diversity. We find pervasive phylogenetic signal in mimicry rings and in color patterns associated within polymorphic species. Only a small set of mimicry rings show high phylogenetic diversity. We identify patterns of saturation in the accumulation of new mimicry rings and in the number of evolutionary convergences towards the most species-rich mimicry rings. Those saturation patterns are likely caused by niche filling along various ecological dimensions, within and among the mimetic communities living in sympatry. The time-dependent effects detected in our study illustrate how neutral processes and ecological interactions interact and shape species and phenotypic diversification.

## Introduction

Identifying how microevolutionary processes and ecological interactions occurring at the community level can affect macroevolutionary dynamics has been a major focus of evolutionary biology (Reznick & Ricklef, 2009, Rolland et al. 2023). Despite a long interest, evidence for an effect of ecological interactions on macroevolutionary patterns of species diversification is scarce. Limitations lie primarily in the difficulty of testing such effects, in particular because species interactions are usually difficult to characterize, and because predicting the effects of processes happening at small spatial and temporal scale on macroevolutionary dynamics is challenging (Rolland et al. 2023). Different ecological interactions (e.g. competition, parasitism, mutualism) may indeed underlie drastically different macroevolutionary trends and involve one or several guilds of species, therefore raising questions about the relative effects of phylogenetic constraints *vs.* selection stemming from these ecological interactions.

Here, we focus on Müllerian mimicry, a widespread and well-documented mutualistic interaction that offers a remarkable opportunity to study the feedbacks between ecological interactions and macroevolutionary processes (Kunte et al. 2021). The evolutionary convergence of warning signals shared by co-occurring defended species, *i.e.* Müllerian mimicry, is one of the most compelling examples of ecological interactions driving trait evolution in multiple distantly related species. Müllerian mimicry among co-occurring species arises through positive frequency-dependent selection exerted by predators that more quickly learn to avoid locally abundant warning signals (Müller 1879, Sherratt 2006). Because the cost of predator learning is diluted among individuals of all the species that share the same warning signal, Müllerian mimicry is a mutualistic interaction; individuals from a given species benefit from co-occurring with individuals belonging to co-mimetic species, thereby promoting wing color pattern convergences in distantly related species. Species that share a common warning color pattern within communities are referred to as mimicry rings.

Müllerian mimicry has evolved independently multiple times in animals, including, among others, frogs (e. g., Symula et al. 2001), snakes (e. g., Sanders et al. 2006), catfish (Alexandrou et al. 2011), butterflies and moths (e. g., Beccaloni 1997), bees and wasps (e. g., Wilson et al. 2012, Ezray et al. 2019) and beetles (e. g., Muñoz-Ramírez et al. 2016). The phenomenon is particularly well-studied in butterflies, in particular in the species-rich neotropical tribe Ithomiini (396 species).

Numerous theoretical, experimental, and field studies have shown that predation is a strong selective force driving the convergent evolution of warning signals such as color patterns (e. g., Sherratt 2006, Elias et al. 2008, Gompert et al. 2011, Willmott et al. 2017, Birskis-Barros et al. 2021). The strength of frequency-dependent selection resulting from predation has been convincingly shown empirically in *Heliconius* (e. g., Mallet and Barton 1989, Kapan 2001, Chouteau et al. 2016) and ithomiine (Willmott et al. 2017) butterflies. Importantly, this selection driven by ecological interactions has led to striking convergence among sympatric species belonging to phylogenetically distant families, such as erebid moths and nymphalid butterflies (Beccaloni 1997). Because mimicry involves strong resemblance across phylogenetically distant species, it provides an excellent context to test the relative importance of ecological interactions and phylogenetic constraints on macroevolutionary dynamics.

Several lines of evidence show that mimicry drives the evolution of multidimensional ecological niches in the species involved, including convergence in multiple traits (e.g. host-plant preference, flight height and flight behaviour, wing morphology, altitudinal and climatic niche) (Willmott and Mallet 2003, Elias et al. 2008, Hill 2010, Gompert et al. 2011, Chazot et al. 2014, Aubier and Elias 2020, Hill 2021, Doré et al. 2023, Page et al. 2024). Mimicry is thus an ecological interaction that plays a key role in the community assembly across multiple sympatric species with various levels of phylogenetic divergence.

In addition to structuring communities, switches in mimetic patterns may act as a driver of speciation in mimetic lineages, since differences in wing patterns among mimetic species drive (Jiggins et al. 2001, Chamberlain et al. 2009, Merrill et al. 2011), and non-mimetic hybrids between taxa belonging to distinct mimicry rings suffer stronger predation (Merrill et al, 2012). Mimetic color patterns are thus often considered as so-called magic traits (*i. e*., traits involved in mate choice that are also under divergent ecological selection, Gavrilets 2004, Servedio et al. 2011), and as one factor contributing to high diversification rates in some clades of mimetic butterflies (Jiggins et al. 2006, Kozak et al. 2015). Mimicry is also predicted to fuel species diversification through the formation of a spatial mosaic of mimicry rings (Aubier et al. 2017), because such a mosaic should allow the persistence of different species occupying similar ecological niches in different patches of the mosaic.

Here, we investigate whether mimetic interactions affect the macroevolutionary dynamics of diversification of mimetic species and their warning signals, as well as the species composition of mimicry rings. We specifically focus on: (1) the diversification dynamics of colour patterns through time and (2) the consequences of such phenotypic diversification on the variation in species richness and phylogenetic diversity among mimicry rings. In order to address these two questions in the absence of a clearly established framework of hypotheses we propose nine predictions to be tested (Figure 1). We used the largest tribe of mimetic butterflies, namely the Ithominii, by combining phylogenetic information with a dataset of color patterns for 339 Ithomiini species (Chazot et al. 2019, Doré et al. 2021, 2022).

**Figure 1.**
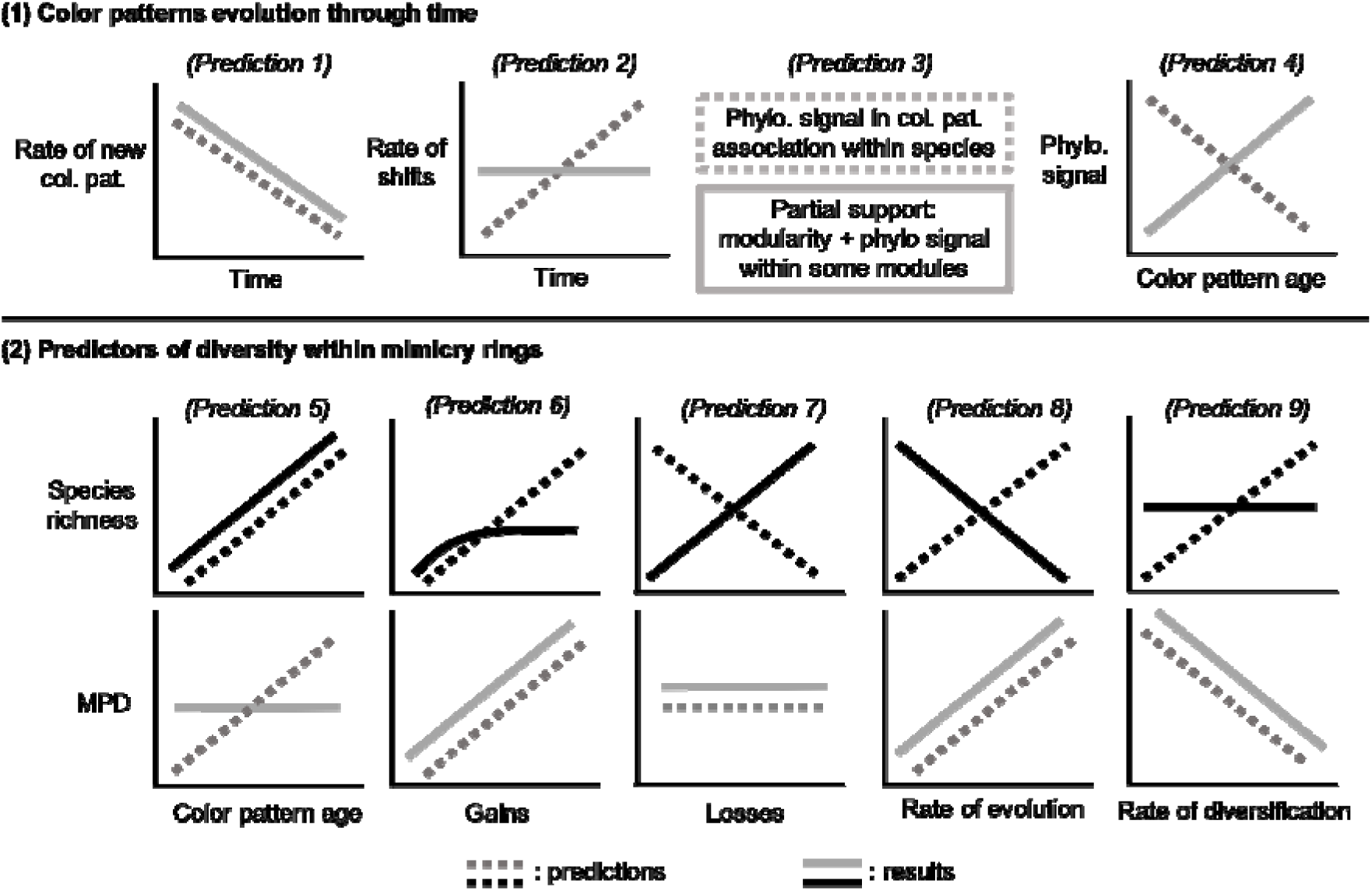
Graphical summary of our nine predictions tested in this study and the results we obtained for Ithomiini butterflies. Predictions 5-9 are subdivided into predictions for species richness and MPD. Predictions are shown with dashed lines and results in plain lines.

## Material and Methods

### Data

We used the time-calibrated phylogeny of Ithomiini from Chazot et al. (2019), which contains 339 species of Ithomiini (85% of species). We used Doré et al (2021)’s color pattern classification and species data, resulting in 44 mimicry rings (Figure 2, Table S1). Mimicry ring species richness in Ithomiini ranges from a single to over 70 species, and, due to local or geographic wing pattern polymorphism, at least half of Ithomiini species belong to two or more mimicry rings.

**Figure 2.**
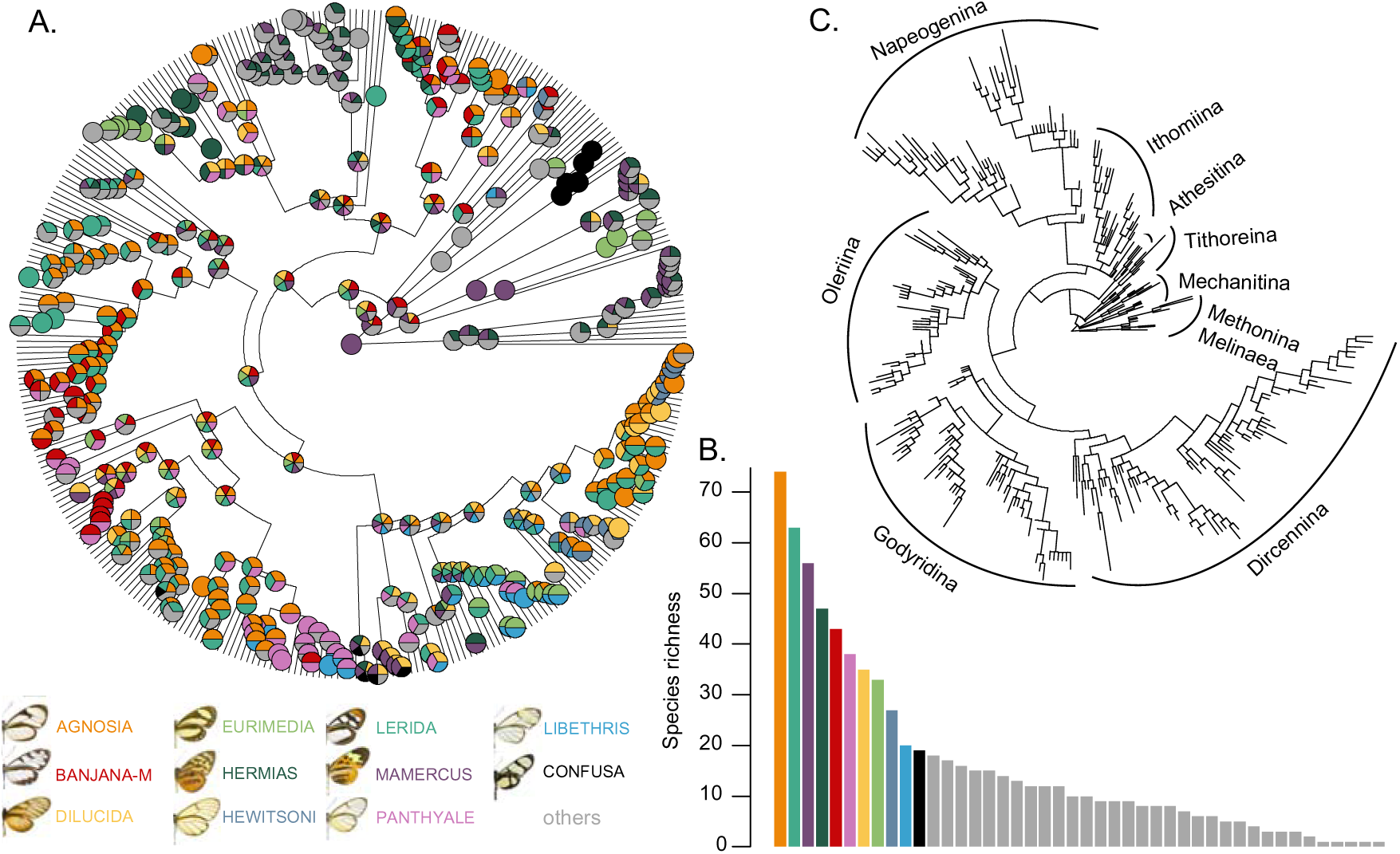
A. Example of a history of color pattern evolution drawn from the posterior distribution of the Bayesian inference. Only the 11 most species-rich mimicry rings are colored. Pie charts do not represent posterior probabilities, rather, they show all mimicry rings that were inferred at a given internal node at this given sample from the posterior distribution. B. Ranked species richness of mimicry rings. C. Phylogenetic tree of Ithomiini with branch lengths rescaled according to the average number of shifts estimated across 100 ancestral histories drawn from the posterior distribution.

#### Ancestral state estimations

In our dataset, 200 out of the 339 species studied show color pattern polymorphism. We therefore estimated ancestral color patterns across the Ithomiini tree using the method described in Braga et al. (2020). This method, initially designed for estimating ancestral host-parasite associations on a phylogenetic tree, allows current and ancestral species to belong to multiple mimicry rings, due to wing color pattern polymorphism. We used the 2-state model and the inference strategy of Braga et al. (2020) as implemented in RevBayes (Höhna et al., 2016) to infer the history of color pattern evolution, based on the color patterns known for each extant species. Model parameters and ancestral states were inferred using Markov Chain Monte Carlo (MCMC). We ran two chains for 8 x 10^4^ iterations, discarding the first 10% as burn-in and sampling every 50 iterations, then confirmed that the chains converged to the same posterior distribution by calculating the Gelman diagnostic (Gelman and Rubin, 1992) in the R package *coda* (Plummer et al., 2006). Results from the MCMC chain with the highest effective sample size are presented. All RevBayes and R scripts are available at github.com/maribraga/Ithomiini_color.

Because polymorphic states can be inferred as ancestral states, the complexity of the evolutionary history estimated greatly increased. We thus took advantage of the Bayesian framework by performing all follow-up analyses on a posterior distribution of evolutionary histories.

### Dynamics of color pattern evolution through time

*Prediction 1: The rate of emergence of new patterns decreases through time*, because the selective advantage of displaying an already prevalent color pattern should favor convergence rather than innovation (Figure 1).

*Prediction 2: The rate of color pattern shifts increases through time as the overall number of color patterns rises*, because an increasing number of color patterns provides new opportunities for convergence (Figure 1).

To test these predictions, we extracted the timing of color pattern evolutionary events (*i.e.*, gains and loss) for each independent history sampled from the posterior distribution of the ancestral state estimations. We extracted the earliest timing of emergence of each color pattern to calculate the cumulative sum of new color patterns through time within 1 My time bins. Using a sliding-window analysis, we also counted the total number of color pattern shifts (irrespective of color pattern identity) occurring within 1 My time bins to estimate the number of shifts through time. Since both statistics are expected to increase proportionally with branch length, we obtained the rate of new pattern emergence (Prediction 1) and the rate of color pattern shift (Prediction 2) for each 1 My time-bin by dividing the number of new patterns and the number of shifts, respectively, by the sum of branch lengths in that bin.

*Prediction 3: There is a significant phylogenetic signal in color pattern associations within species*, caused by phylogenetically conserved developmental pathways of color patterns and/or parallel co-evolution of color patterns across species, since polymorphic species can simultaneously shift to different mimicry rings.

We evaluated whether polymorphic species exhibit random color pattern associations or instead, whether they represent repeated association of the same color patterns. We used statistical tools designed for ecological network analyses to test for modularity in the network of color pattern associations, where modules represent clusters of color patterns frequently associated in polymorphic species. We used the Beckett (2016) algorithm to identify the network configuration maximizing the proportion of interactions within modules. We repeated the module identification 100 times and calculated the frequency of pairwise associations of color patterns within modules. We used the package *qgraph* to build the network of pairwise associations.

We then tested whether the species composition of each module was phylogenetically random or instead phylogenetically structured. We measured phylogenetic diversity in the modules by comparing the mean phylogenetic distance (MPD) between species belonging to the same module to the distribution of mean phylogenetic distances after permutations of color patterns between modules. High MPD score indicates a module comprising more distantly related species than expected by random community assemblage, *i.e.* resulting from color pattern convergence between distantly related lineages. Conversely, a low MPD indicates a module shared by species more closely related than a random combination of species. MPD values falling into the 95% interval of the distribution obtained from the permutations indicate no departure from random evolution of color pattern, *i.e.* driven neither by convergence events nor by phylogenetic proximity. Since modules are not identical in different iterations, we cannot use the entire posterior distribution for this test. We performed the MPD test on one iteration of the module search algorithm.

*Prediction 4: Phylogenetic signal of color patterns decreases with the age of the color pattern* (Figure 1). Positive frequency-dependent selection on color patterns is expected to drive convergence among distantly related lineages, thereby decreasing phylogenetic signal. Greater time since the emergence of a color pattern allows for color pattern convergence involving more distantly related species (Kunte et al. 2021).

To assess the phylogenetic signal of color patterns, we computed the D statistics (Fritz & Purvis 2010) for each mimicry ring (while merging all other mimicry rings) with more than 3 species, with permutations to identify significantly conserved and overdispersed mimicry rings. A D value of 0 indicates Brownian motion evolution across the tree, and a value of 1 random evolution uncorrelated with phylogeny. D<0 indicates phylogenetic clustering (i. e., conservatism) while D>1 indicates phylogenetic overdispersion.

We obtained the age of color patterns by extracting the timing of color pattern gain and loss events for each independent history sampled from the posterior distribution of the ancestral state estimation analysis. The oldest timing of emergence of a color pattern, in absolute value, was considered as the age of the color pattern. To test our prediction, we fitted a linear model with D as response variable, and color pattern age as explanatory variable.

### Species richness and phylogenetic diversity within mimicry rings

In Ithomiini butterflies there is a huge disparity in mimicry ring species richness (i. e., the number of species in a mimicry ring), which ranges from a single species to over 70 (Figure 2). We estimated mimicry ring phylogenetic diversity using an MPD test similar to the tests performed within modules. We compared the mean phylogenetic distance between species belonging to the same mimicry ring to the distribution of mean phylogenetic distances after permutations. Five color patterns were unique to a single species and therefore we did not perform the MPD test for the mimicry rings harboring these color patterns. MPD also strongly varied across mimicry rings, from phylogenetically overdispersed to highly clustered mimicry rings. A linear regression confirmed that MPD does not predict species richness (F= 2.20, Df=37, p=0.146, R2=0.03, Supplementary Figure S2A). To explain such disparity in both species richness and phylogenetic diversity, we estimated the contribution of four processes by which variation in species richness or phylogenetic diversity between mimicry rings may arise (Figure 1).

*Prediction 5: Species richness and phylogenetic diversity of a mimicry ring increase with the age of the associated color pattern* (Figure 1). As a null hypothesis, species richness should increase with time through the accumulation of either speciation without color pattern change or speciation driven by convergence of color pattern. Greater time since the emergence of a color pattern allows for color pattern convergence involving more distantly related species, hence increasing phylogenetic diversity (Kunte et al. 2021).

*Prediction 6*: *Species richness and phylogenetic diversity increase with the number of evolutionary convergence (*Figure 1). Differences in evolutionary gains among color patterns may be observed, either (1) because they confer different advantages such as a higher protection against predation (e. g., because they are locally abundant, or more memorable) or benefits unrelated to aposematism (e.g., thermoregulation, crypsis), or (2) because fewer mutations are required for this pattern to emerge from an ancestral one, for example when the capacity to produce certain pigments and pattern elements is already present as an ancestral state.

We summed the total number of species gain events for each mimicry ring across the posterior distribution of ancestral state scenarios, similar to the calculation of the rate of shifts through time. However, even under stochastic evolutionary processes, the number of gains increases with time. Thus, we divided the gains by the age of the color pattern to obtain a rate.

*Prediction 7*: *Species richness decreases with the number of evolutionary losses of color patterns, while phylogenetic diversity should not be affected if losses are randomly distributed across lineages* (Figure 1). Here, evolutionary losses do not imply lineage extinction but rather evolutionary shifts in color patterns leading to the loss of an ancestral color pattern. We calculated a rate of color pattern loss in the same way we calculated the rate of gain.

*Prediction 8*: *Species richness and phylogenetic diversity of a mimicry ring increase if lineages belonging to that mimicry ring exhibit a higher rate of color pattern evolution* (Figure 1). Such lineages may lead to a high number of convergence events towards the same color pattern, thereby increasing species richness and phylogenetic diversity via convergence and subsequent diversification. While Prediction 6 focuses on tree-wide association between rates of gains and losses, Prediction 8 tries to account for clade-specific rates of evolution that may lead to higher species richness.

We estimated a tip rate of color pattern evolution – species-specific rate of evolution – by first calculating the number of shifts occurring along each branch in the tree across the posterior distribution of ancestral state estimations. The length of each branch of the phylogenetic tree was then rescaled to this number of events. We calculated for every tip in the tree the total rescaled branch length connecting the root to the tips. Because the number of possible evolutionary changes is expected to increase with the number of nodes connecting the root and the tips, we divided the sum of branch length of each tip by the number of connecting nodes to obtain the tip rate of color pattern evolution. Finally, we calculated the median tip rate of evolution for each color pattern.

*Prediction 9: Species richness of a mimicry ring increases, but phylogenetic diversity decreases, with the rate of diversification of the lineages belonging to it* (Figure 1). Although the reasons why a given color pattern could directly affect the probability of a speciation event (without color pattern shift) are unclear, it may be indirectly associated with higher diversification through correlation with other phenotypical, ecological or geographical traits e.g., habitats that favor speciation or limit extinction, such as high-altitude Andean forests (Chazot et al. 2016, 2018, 2019). Higher speciation rate within a mimicry ring will lead to a larger number of species harboring the same color pattern, thereby increasing the species richness of the mimicry ring, while decreasing its mean phylogenetic diversity because most species joining that mimicry ring will be closely related.

To test Prediction 9, we calculated the Diversification Rate (DR) statistics (Jetz et al., 2012) for each tip in the tree. DR is a tip rate of diversification calculated as the inverse of the equal-split measure (Redding & Mooers 2006). We calculated the median DR for each mimicry ring from all the species belonging to the same mimicry ring. We used MPD of mimicry rings as before.

### Statistical analyses

We tested for relationships between species richness or phylogenetic diversity and each explanatory variable using linear regressions. However, the age of color patterns correlated with many of our explanatory variables. In particular, both species richness and the rate of shifts increased with age. Therefore, we also calculated the residuals of the linear model between species richness and color pattern age, then repeated the statistical analyses using those residuals as the dependent variable. We found no relationship between MPD and color pattern age and thus did not perform the test for residual MPD. All statistical tests were performed using the median predictor scores calculated over 100 posterior evolutionary histories.

## Results

### Color pattern evolution through time

The number of color pattern shifts increased over time, but the rate of shifts remained nearly stable (Figure 3A and 3C), providing no support for our prediction that an increasing number of colour patterns fuels colour pattern shifts (Prediction 2). The rate of new pattern emergence decreases through time, moderately until about 15 My ago, and more abruptly after that, with hardly any new pattern appearing during the last 5 My (Figure 3B and 3D), consistent with saturation of colour patterns (in line with Prediction 1).

**Figure 3.**
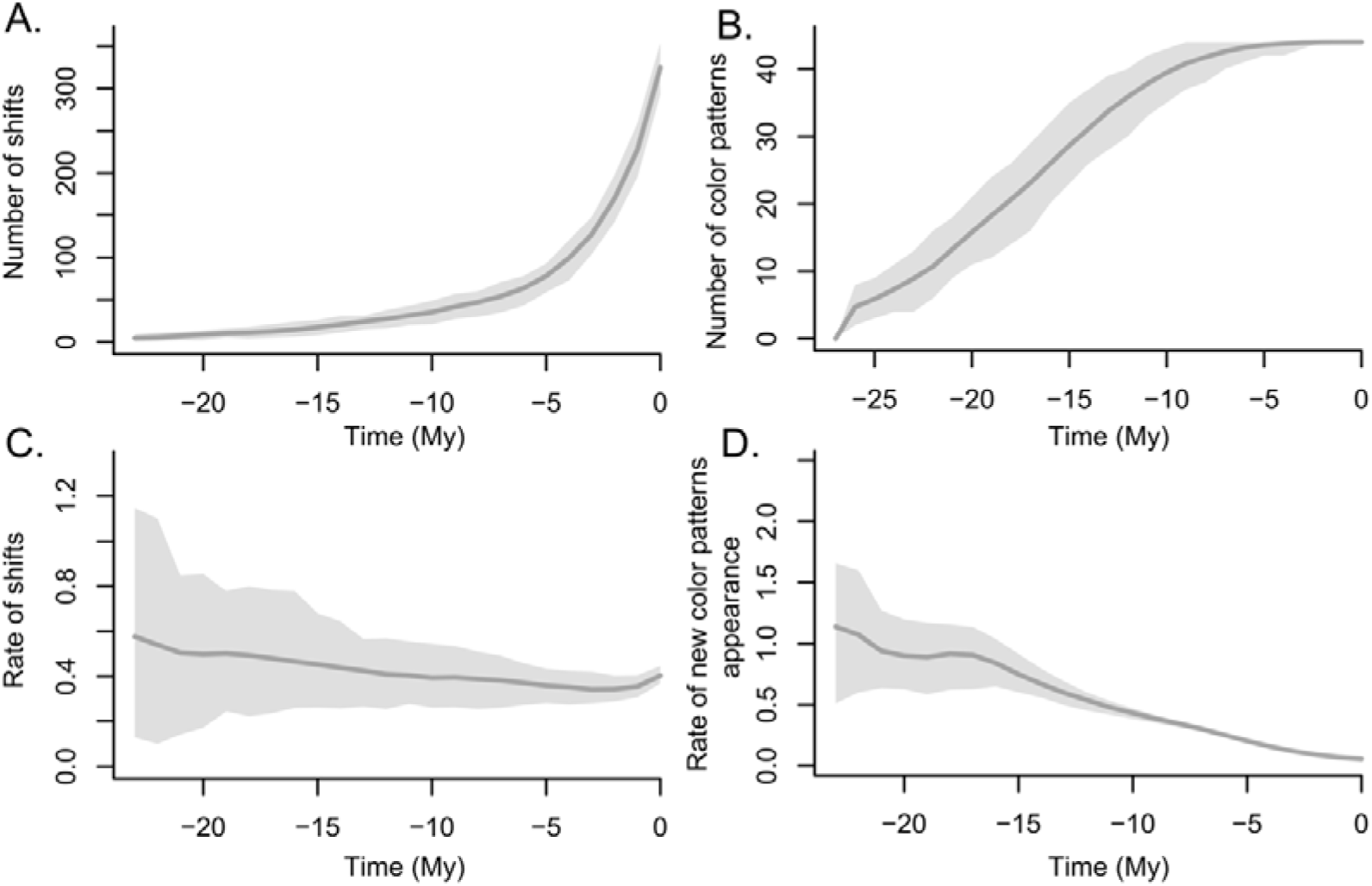
Number and rate of color pattern shifts through time. Cumulative number of new color patterns through time and rate of new color pattern appearance through time. Grey polygons show the 95% confidence interval from 100 posterior ancestral state estimations, line showing the median.

### Patterns of species richness and modularity

There was high heterogeneity in extant species richness among mimicry rings, with a mean of 16.3 species per mimicry ring, ranging from 74 species sharing the same warning signal (color pattern AGNOSIA), to a single species (ACRISIONE, DERCYLLIDAS, HUMBOLDT, PRAESTANS, VESTILLA) (Figure 2). We found modularity in the network of interactions, indicating that certain combinations of color patterns within polymorphic species occur more often than expected at random. A single iteration of the Beckett (2016) algorithm identified nine modules (Figure 4A). The most species-rich mimicry rings were not concentrated in a single module. For example, the six most species-rich mimicry rings were distributed across four different modules. We found significantly low Mean Phylogenetic Distance (MPD) scores for four of these modules (module 2, 3, 4, 9, Figure 4A). A single module had significantly high MPD (module 6). This module contained most of the patterns with black, orange and yellow opaque colors (often referred to as ‘tiger’ color patterns). The generally low MPD score suggests that the association of different colour patterns within species partially stems from phylogenetic proximity between species, therefore providing partial support for Prediction 3. Nevertheless, not all modules were supported based on 100 iterations of the Beckett algorithm (Figure 5). Module 2, 4 and 6 were well supported and module 3 and module 1 only partly supported (Figure 5). This analysis of the strength of color patterns associations also clearly highlights a few color patterns generally not involved in polymorphism, such as LIBETHRIS and DUILLIA. By contrast, DOTO appears to be involved in many different iterations of polymorphism.

**Figure 4.**
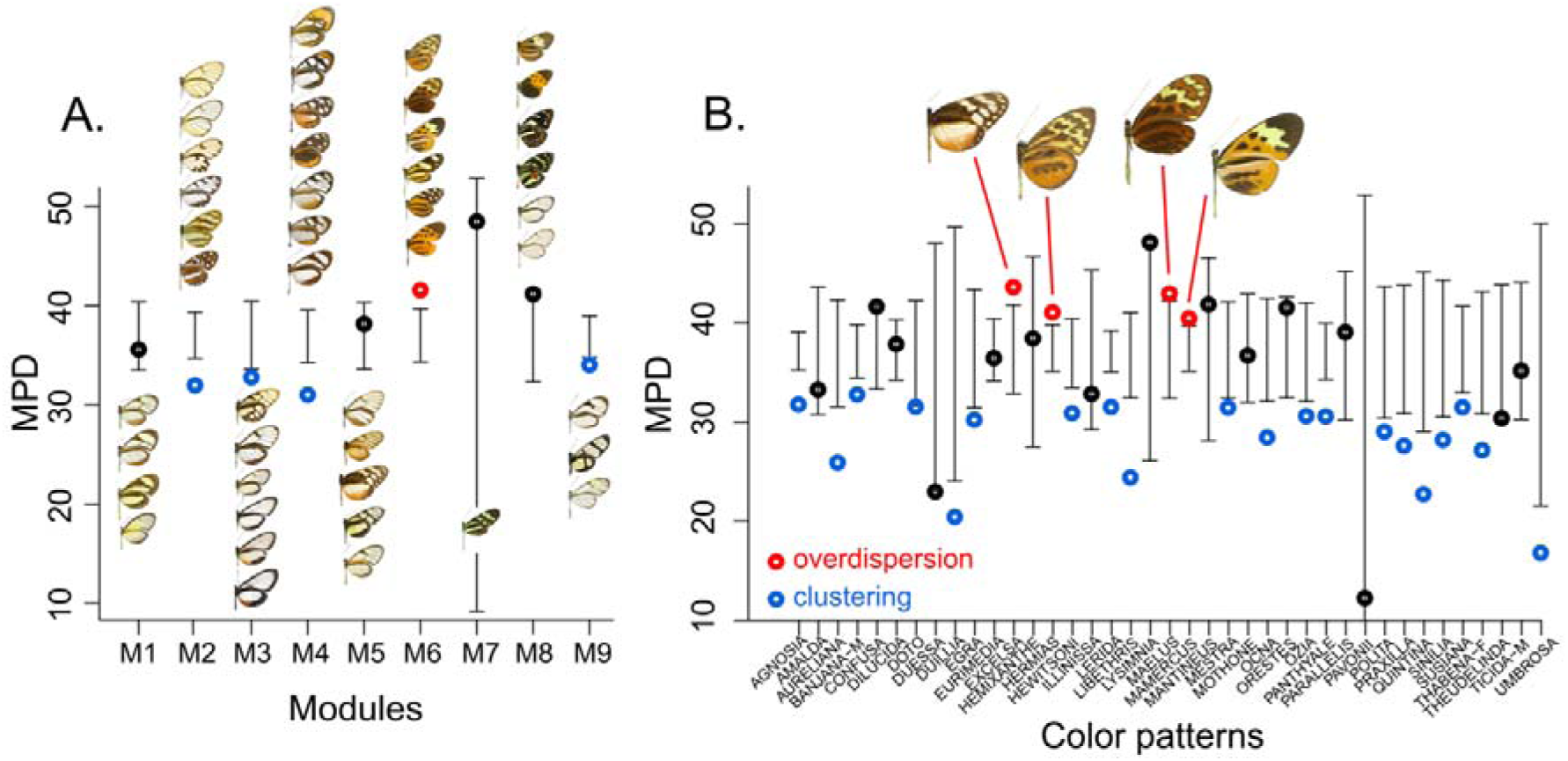
Phylogenetic diversity within modules (A.) and within mimicry rings (B.). Circles indicate the observed mean phylogenetic distance (MPD) score. Bars indicate the 95% confidence interval of MPD scores after 1000 permutations. Butterfly images show the color pattern associated with each mimicry ring or module.

**Figure 5.**
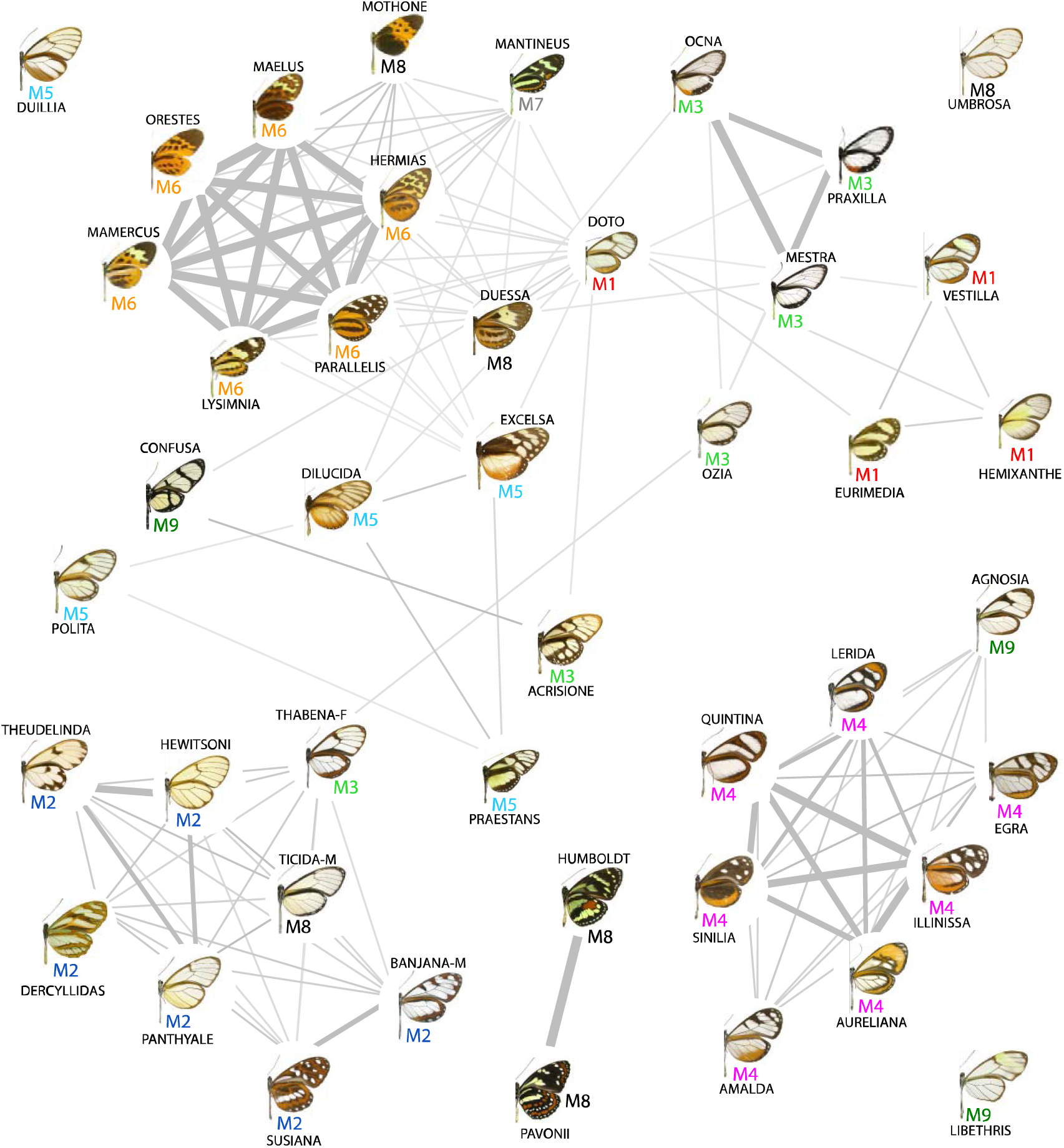
Graphical representation of the frequencies of pairwise associations of color patterns within polymorphic species calculated from 100 iterations of the Beckett (2016) algorithm. Only pairwise associations with a frequency higher than 0.25 are shown. Color pattern names are provided with each picture. Labels M1-M9 correspond to the module assignments as provided in Figure 4.

### Phylogenetic signal within mimicry rings

Using the D statistics, we found significant phylogenetic signal for 14 color patterns and overdispersion for 12 mimicry rings (Figure 6A). The D statistics increased as color patterns emerged more recently, indicating higher phylogenetic signal in older mimicry rings, *i.e.* opposite to Prediction 4 (Figure 6A).

**Figure 6.**
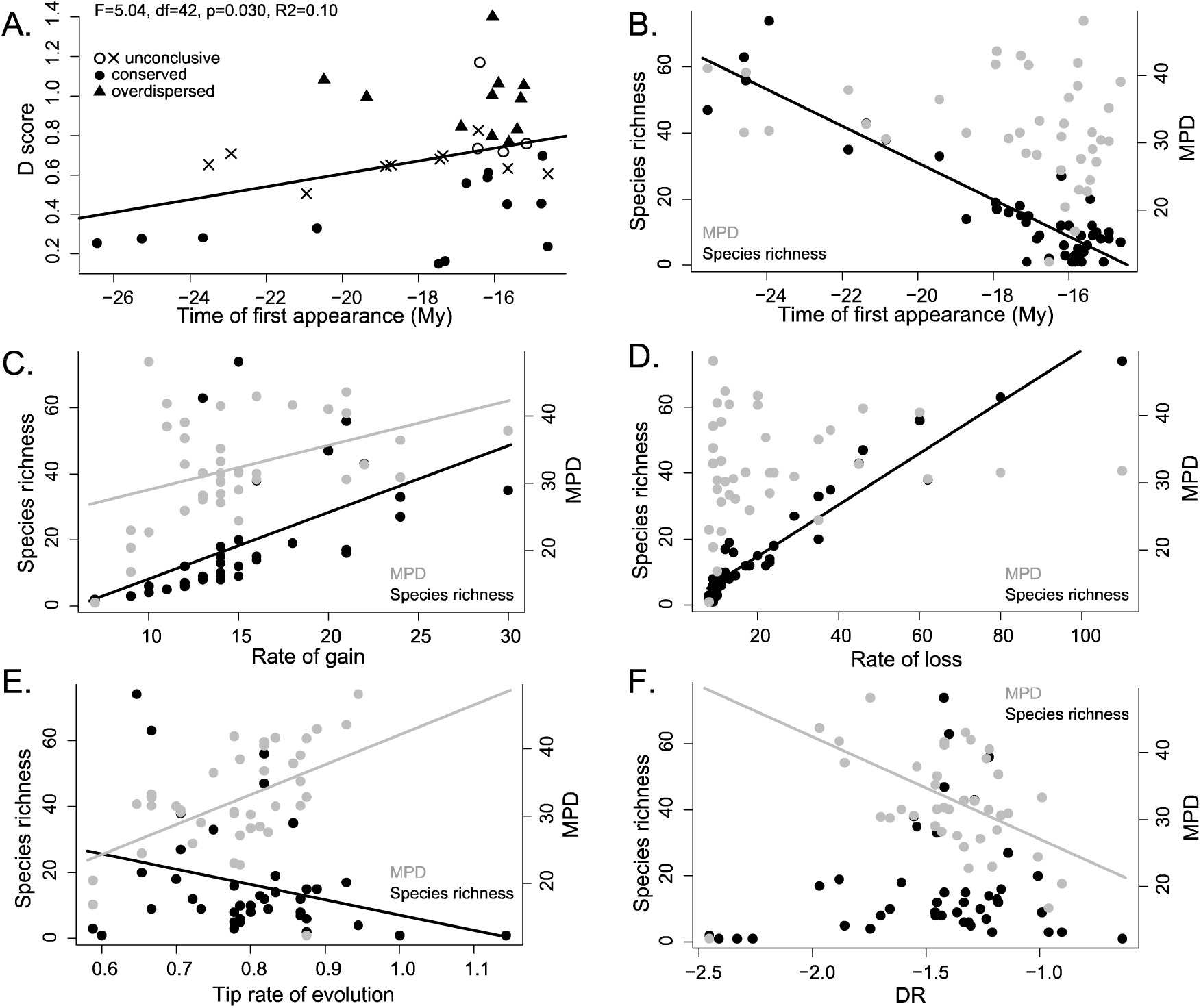
Predictors of mimicry ring species richness and phylogenetic diversity. (A.) Relationship between phylogenetic signal for each color pattern (D score) and the time of first appearance. (B. - F.) Species and MPD as functions of color pattern age (B.), rate of gain (C.), rate of loss (D.), tip rate of color pattern evolution (E.), diversification rate (F.). Black dots correspond to species richness, gray dots correspond to MPD. The linear models fitted for each predictor individually are shown when each variable significantly predicted either mimicry species richness or MPD. Results of the regression analyses performed are provided in Table 1.

**Table 1.**
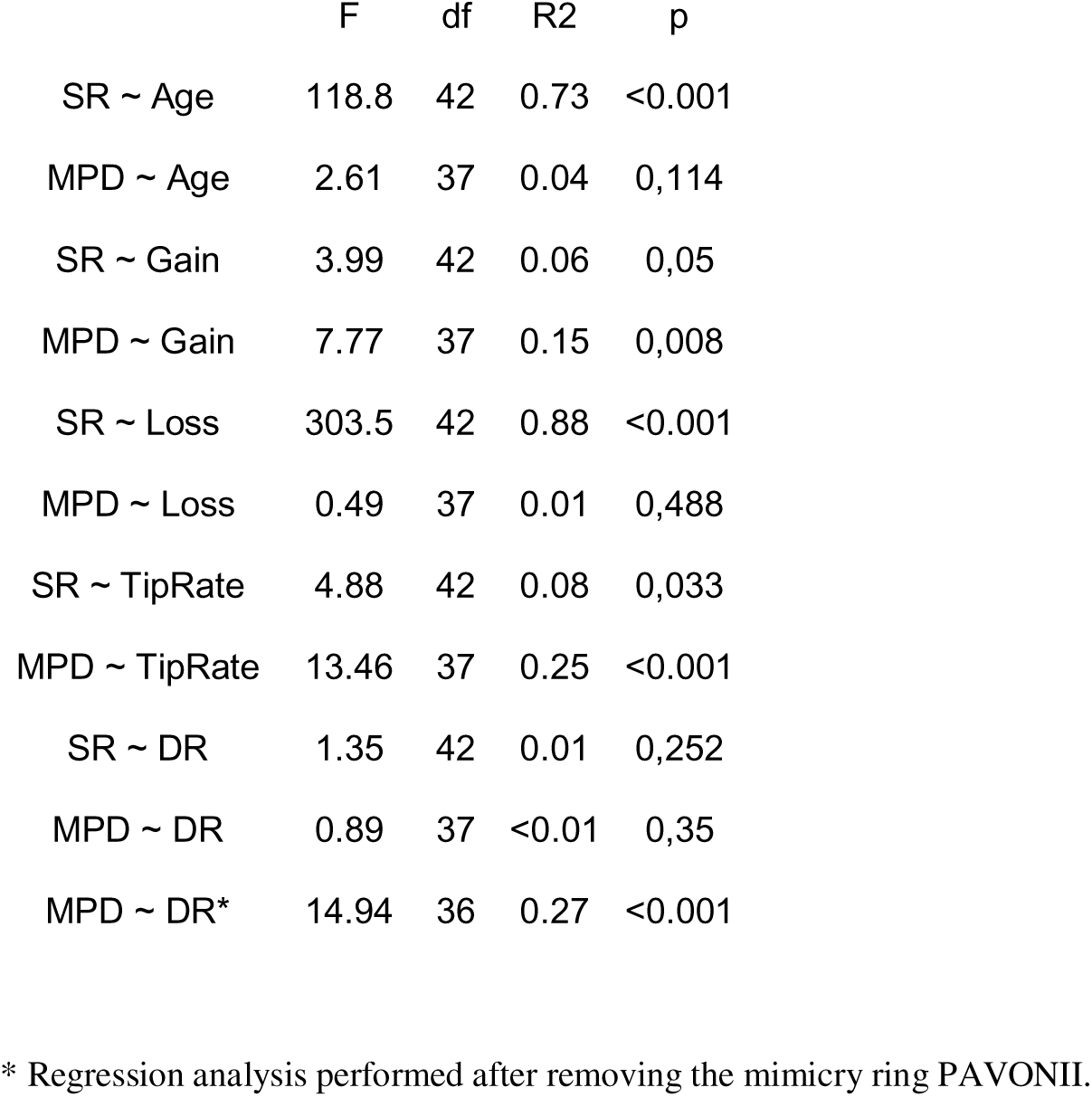
Results of the regression analyses performed between either mimicry species richness or MPD and five predictors of diversity: age, rate of gain, rate of loss, tip rate of color pattern evolution, diversification rate (DR).

### Predictors of mimicry ring species richness and phylogenetic diversity

Using the MPD test, we found significantly low mean phylogenetic diversity within 20 mimicry rings (Figure 4B). By contrast, only four mimicry rings showed significantly high mean phylogenetic diversity (EXCELSA, HERMIAS, MAELUS, MAMERCUS, ‘tiger’ color patterns with black, orange and yellow opaque colors).

We found that the age of color patterns (Prediction 5), the rate of color pattern losses (Prediction 7) and the tip rate of evolution (Prediction 8) were the main predictors of mimicry ring species richness (Figure 1, 6, Table 1), but the observed patterns for Predictions 5 and 7 are opposite to those expected (Figure 6D&E). In addition, the smallest mimicry rings tended to harbor the lowest gain rates, but the relationship between the rate of gains and species richness was weak (p = 0.05, Figure 6C, Table 1, Supplementary Figure S2). Diversification rate (Prediction 9) seemed higher in species-rich mimicry rings but the relationship was not significant (Figure 6F, Table 1). Overall, species-rich mimicry rings were older, experienced higher rates of color pattern loss and assemblages of lineages with high rates of evolution (Figure 1). When performed on richness-age residuals, the rate of gains, losses and tip rate of evolution significantly predicted species richness (Supplementary Figure S2).

When testing the same predictors for phylogenetic diversity we found that neither color pattern age (Prediction 5) nor the rate of losses (Prediction 7) predicted MPD (Figure 1, 6, Table 1, Supplementary Figure S2). However, we found that mimicry rings with higher phylogenetic diversity had higher rates of gain (Prediction 6) and higher tip rate of color pattern evolution (Prediction 8, Figure 1, 6, Table 1). We initially found no relationship between MPD and DR. However, we identified one outlier in our analysis (mimicry ring PAVONII shared by only two species). When we performed the regression test again without PAVONII, we found that high MPD mimicry rings were associated with low DR (Prediction 9, Figure 6, Table 1).

## Discussion

### Strong phylogenetic signal and convergences in the formation of mimicry rings

Despite selection by predation favoring convergence among distantly-related species, we found significant phylogenetic signal in 14 mimicry rings. Modularity in the network of mimetic interactions also indicates that there is a non-random co-occurrence of color patterns in polymorphic species. Out of the nine detected modules, half were phylogenetically clustered (Figure 4), although the interpretation of this result is limited by the poor support we found for some of these modules. Furthermore, polymorphic states at nodes were generally estimated from our analyses. This suggests that phylogenetic inheritance is not limited to single color patterns but also frequently affects the ability to produce sets of different patterns, maybe generated by similar developmental pathways. It is also possible that ecological similarities between co-mimetic species lead to repeated co-variation of color patterns when these co-mimetic species co-occur over large geographical scale.

We predicted that phylogenetic signal should decrease with color pattern age (Prediction 4, Kunte et al. 2021) but we found the opposite result (Figure 1, 6). Even though the number of convergence events also increases with time (in line with Prediction 4), it is probably outweighed by speciation without color pattern changes and constrained by the time-dependent effects as discussed below.

Mean phylogenetic diversity was significantly higher than expected under a random distribution of the color patterns within four mimicry rings (EXCELSA, HERMIAS, MAELUS, MAMERCUS). All of them but EXCELSA are part of the same module, the only module also phylogenetically overdispersed (Figure 4). While wing transparency is very common in Ithomiini, with most of the mimicry rings having wings with transparent patches, the four highly convergent color patterns mentioned above are all non-transparent and are frequently found outside of Ithomiini, such as in heliconiine butterflies (Nymphalidae: Heliconiini), danaines (Nymphalidae: Danainae: Danaini) or in pericopine tiger-moths (Erebidae: Pericopina). The elevated number of distantly-related species displaying these color patterns, both within and outside Ithomiini butterflies, may stem from several developmental or selective factors: 1 - the retention of genetic abilities that could be ancestral to all Lepidoptera (phylogenetic signal hypothesis), 2 - the fact that these patterns can be produced by a variety of genetic toolkits, thus making them more likely to appear independently (evolutionary convergence hypothesis), 3 – opaque patterns being more conspicuous and therefore more easily learnt and remembered by predators (natural selection hypothesis).

### Color pattern evolution through time: evidence for ecological saturation?

We initially predicted that the frequency of color pattern shifts would increase through time as a result of the increasing number of possibilities for convergence provided by the increasing number of color patterns with time (Prediction 2, Figure 1). Yet, across all color patterns, we found hardly any variation in the dynamics of shifts through time (when accounting for the increasing number of lineages through time, Figure 3). Therefore, the increasing complexity of the network of interactions over time did not result in an increase in the overall rate of evolutionary shifts. The fact that the overall rate of shifts remained stable while the number of patterns increased implies that the rate of gains and losses per pattern decreased through time.

In parallel, we found a strong dampening in the rate of appearance of new color patterns, with very few new patterns appearing over the last 10 My, suggesting the existence of a large-scale upper limit to the number of color patterns. Such dampening could stem from a mimicry saturation effect. While novel color patterns tend to be eliminated by positive frequency-dependent selection, they may still become established due to drift followed by selection (shifting balance, Mallet and Joron 1999). Evolution of such novel color patterns is especially favored when selection from predation is relaxed (Chouteau and Angers 2012), as may be the case in habitats devoid of an established mimetic community (Sherratt 2006). Therefore, mimicry diversity can increase as long as there are empty ecological niches (e. g., new geographical regions, new elevations, new microhabitats), with communities of naive predators, to colonize (Aubier et al. 2017). When all niches have been colonized by mimetic butterflies, even if there is still room to accommodate more species because resources are not yet limiting, we can expect a strong filtering on color patterns, which are under selection to match the local mimetic community (Kunte et al. 2021, Aubier et al. 2017, Sherratt 2006, Mallet & Barton 1989). In Ithomiini, all geographical regions and likely most altitudinal niches were already colonized five million years ago, but the number of ithomiine species at that time was only about 25% of the number of extant species (Chazot et al 2019). Such conditions likely provided opportunities for further species accumulation in those different regions and habitats, either through speciation or colonization (Chazot et al. 2016, 2018). Nevertheless, mimetic communities may have been well established already, thereby favoring the establishment of species with an already existing color pattern locally. As species continued to be formed, novel color patterns were likely selected against.

We also found evidence of saturation in the balance between the rates of color pattern gains and losses (Supplementary Figure S2). The rate of losses increased with the age of mimicry rings and in species-rich mimicry rings (opposite to our Prediction 6). However, while the rate of losses increased linearly with species richness and equaled the rate of gains for recent/species-poor mimicry rings, the rate of gains in the oldest/species-rich mimicry rings were disproportionately low (Supplementary Figure S2). Overall, for mimicry ring species richness to increase over time despite a high rate of evolutionary losses, the rate of speciation without loss of ancestral color pattern must exceed the rate of color pattern loss. This result further supports the retention of color patterns over evolutionary time. It is also consistent with the significant decrease of tip rate of color pattern evolution in species-rich mimicry rings (in a direction opposite to our Prediction 7). The rate of color pattern evolution declined in lineages belonging to species-rich mimicry rings, and the decline in the rate of gains highlighted above is probably responsible for that.

The dampening of the rate of evolutionary gains is also a plausible explanation for the absence of a relationship between MPD and color pattern age. We predicted that MPD should increase with color pattern age via the accumulation of convergence events. We indeed found that the rate of evolutionary gains predicts MPD. But the time-dependency of this rate of evolution probably constrains the accumulation of phylogenetic diversity as mimicry rings accumulate species.

Altogether, our results point at a saturation effect within mimicry rings, which could be due to resource limitation; co-mimetic species tend to share habitats (Chazot et al. 2014, Doré et al. 2023), microhabitats (Elias et al. 2008, Willmott et al. 2017) and even larval hostplants (Wilmott and Mallet 2003). After a mimicry ring becomes too species-rich the cost of competition among co-mimics may outweigh the anti-predator benefits of joining a very large mimicry ring.

### The age of mimicry rings predicts species richness but not phylogenetic diversity

In our results, time emerged as a major factor determining mimicry ring species richness, which altogether generates a saturation effect limiting new evolutionary convergence. We found that mean phylogenetic diversity of mimicry rings is determined by different processes than species richness. In agreement with our predictions, mean phylogenetic diversity was high in mimicry rings where the color pattern is frequently acquired and whose lineages have high rates of color pattern evolution. But phylogenetic diversity of a mimicry ring results from the balance between diversification without color pattern shift and the rate of evolutionary convergence and we found that MPD decreased with high diversification rates. This result further supports the relative importance of color pattern retention during diversification. Even though evolutionary shifts increased over time and MPD increased with the number of evolutionary shifts, color pattern age did not predict MPD. This is likely another side effect of the dampening of convergence events in species rich mimicry rings that indirectly bound phylogenetic diversity in species rich mimicry rings.

## Conclusion

Mimicry is a remarkable example of an interaction evolving through natural selection. How such selection translates into the dynamics of phenotypic evolution and species accumulation at macroevolutionary scale has previously been poorly studied (but see Aubier et al. 2017). Our results on Ithomiini support mechanisms of diversification that are independent from mimicry. Time alone is a strong predictor of the number of evolutionary events and species richness in mimicry rings. We also found evidence of phylogenetic signal during diversification, which directly influences the relative importance of speciation without color pattern change over convergence in determining mimicry ring composition. Therefore, the network of interactions and the composition of mimicry rings appear to be driven in a large part by processes not associated with mimicry.

Nevertheless, we find evidence that ecological saturation possibly linked to mimetic interactions did affect the dynamics of color pattern evolution and species diversification. This conclusion comes from several lines of evidence: the strong decrease in the rate of new color pattern appearance, the decrease in color pattern gains relative to losses in species-rich mimicry rings, and the increasing rate of losses in species-rich mimicry rings. Ecological limitations to the overall diversity of color patterns and to the number of species interacting within a mimicry ring seem likely to change the dynamics and directionality of color pattern evolution, and thereby the species composition of mimicry rings.

Our results also have implications for the global biogeography of species interactions. Large networks of mimetic interactions seem to be primarily found at lower latitudes (although there has not been any formal validation of this observation). Our study shows that time is key to building a network of mimetic interactions, both in terms of color pattern diversification and species composition of mimicry rings. Thus, our results support the idea that complex networks of interactions are built over long geological times, such that specific conditions such as the age of biomes or their climatic stability (Fischer 1960) may be key to understanding latitudinal gradients in biotic interactions.

