## Supplementary material for "Bending the course of evolution: how mutualistic interactions affect macroevolutionary dynamics of diversification in mimetic butterflies"

Table S1. Ithomiini_mimicry_data_mtx.nex (separate file) Mimicry ring assignment of Ithomiine species in nexus format. Mimicry rings are labelled in the following order: ACRISIONE, AGNOSIA, AMALDA, AURELIANA, BANJANA-M, CONFUSA, DERCYLLIDAS, DILUCIDA, DOTO, DUESSA, DUILLIA, EGRA, EURIMEDIA, EXCELSA, HEMIXANTHE, HERMIAS, HEWITSONI, HUMBOLDT, ILLINISSA, LERIDA, LIBETHRIS, LYSIMNIA, MAELUS, MAMERCUS, MANTINEUS, MESTRA, MOTHONE, OCNA, ORESTES, OZIA, PANTHYALE, PARALLELIS, PAVONII, POLITA, PRAESTANS, PRAXILLA, QUINTINA, SINILIA, SUSIANA, THABENA-F, THEUDELINDA, TICIDA-M, UMBROSA, VESTILLA.

Supplementary Figure S1. Posterior distribution summary of ancestral state estimations. Color patterns with more than 90% marginal posterior probability shown at nodes, only for the 10 most species rich mimicry rings.

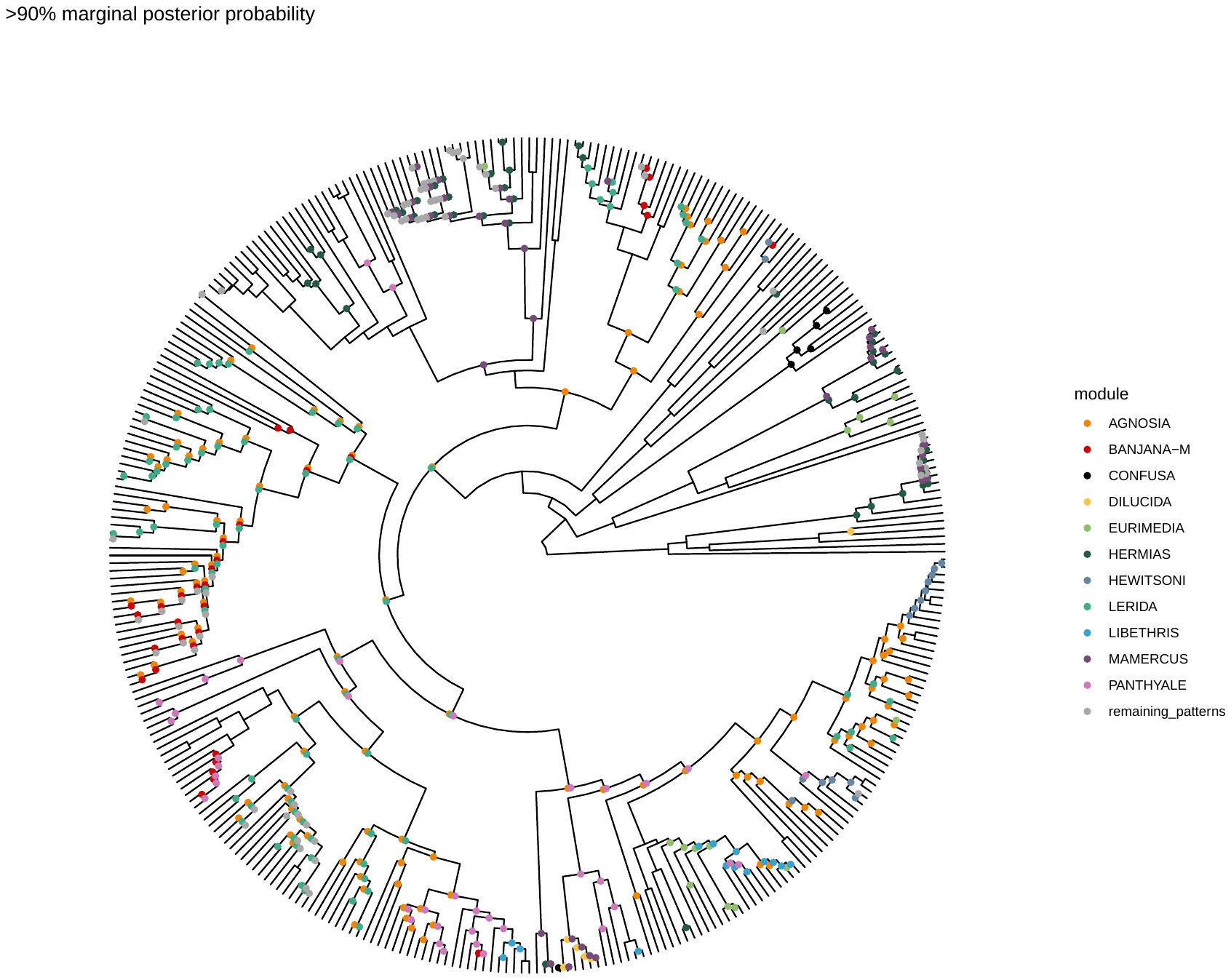

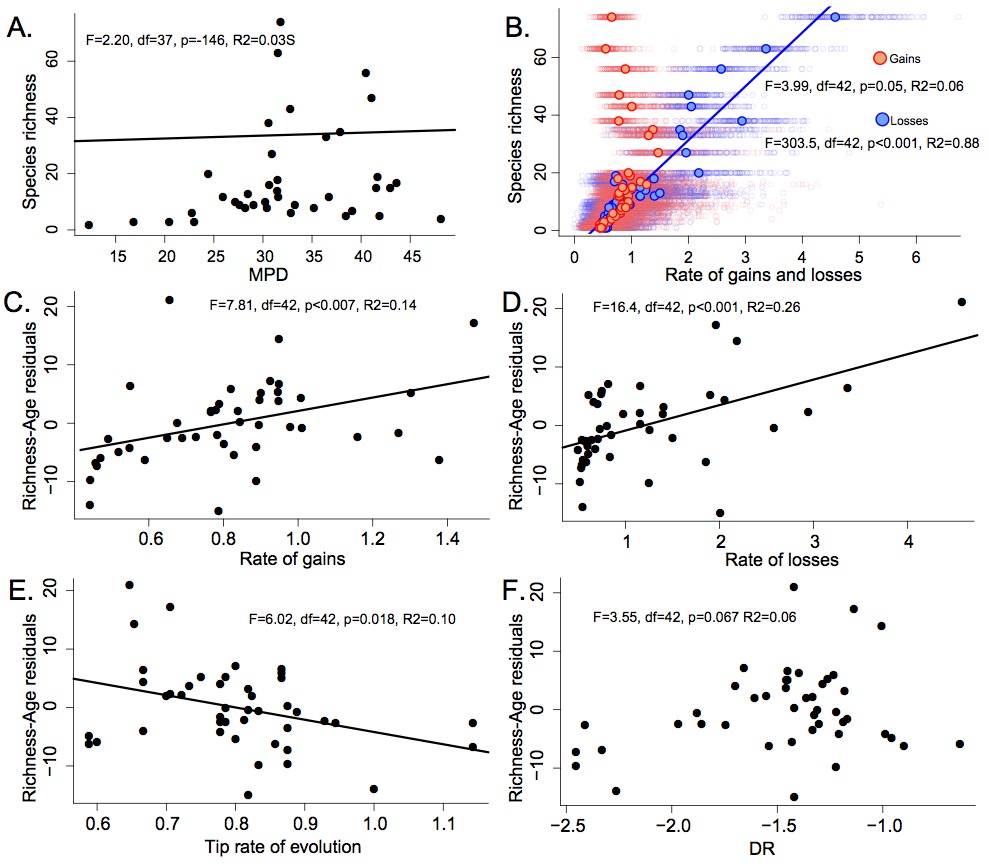

Supplementary Figure S2. A. Mimicry ring species richness as a function of MPD. B. Mimicry ring species richness as a function of the rates of gain and loss. C. Species richness - color pattern age residuals as a function of the rate of gain. D. Species richness - color pattern age residuals as a function of the rate of loss. E. Species richness - color pattern age residuals as a function of the tip rate of evolution. F. Species richness - color pattern age residuals as a function of the diversification rate DR. The results of the regression tests are associated with each graph.
